# Roadkill is a crucial factor in the population decline of migratory monarch butterflies

**DOI:** 10.1101/2024.09.27.615542

**Authors:** Iman Momeni-Dehaghi, Lenore Fahrig, Greg W. Mitchell, Trina Rytwinski, Jeffrey O. Hanson, Joseph R. Bennett

**Affiliations:** Department of Biology, Carleton University; Ottawa, ON, Canada; Wildlife Research Division, National Wildlife Research Centre, Environment and Climate Change Canada; Ottawa, ON, Canada

**Keywords:** Least cost path, migration, migratory monarchs, roadkill, road ecology

## Abstract

The charismatic migratory monarch butterfly population has declined dramatically, likely precipitated by loss of its breeding host plants (milkweed). Whether restoring milkweed would allow monarch recovery depends on whether additional factors currently limit the population. We investigated road mortality as one such factor. Monarchs cross thousands of roads during fall migration, and traffic volume has increased sharply while the population has plummeted. Using estimates of pre-migration distribution, flight patterns, and road traffic, we estimate that at least 61% (range 61% to 99.99%) of migrating monarchs are road-killed each fall. Although there is high uncertainty in our estimate, its magnitude suggests that roadkill could inhibit recovery of the population. Recovery planning should not only consider increasing the monarch’s host plants, but must also address the reality of roadkill.

## Introduction

Monarch butterflies (*Danaus plexippus*) in eastern North America are well-known for their remarkable migration. Every year around mid-August, monarchs start their fall migration from their breeding areas in the USA and Canada toward their overwintering areas in central Mexico. Then, in early spring, the same individual monarchs who migrated to Mexico, migrate back to the southern USA where they establish the first generation of breeding monarchs. Offspring of these monarchs later repopulate the breeding range of monarchs in Canada and the USA, before the fourth and fifth generations initiate the fall migration again in August. The fall migration, completed by a single generation of monarchs, can be more than 4,000 km, one of the longest distances that insects migrate.

The population size of migratory eastern monarchs has declined dramatically since the first population estimate in 1994 (Fig. 1). In 2024, the population was around 73.6% lower than the average population size from 1994 to 2023 (1). This decline has prompted high-level international efforts, such as the Trinational Monarch Conservation Science Partnership (2) aimed at enhancing our understanding of monarch population dynamics and the threats they face.

**Figure 1:**
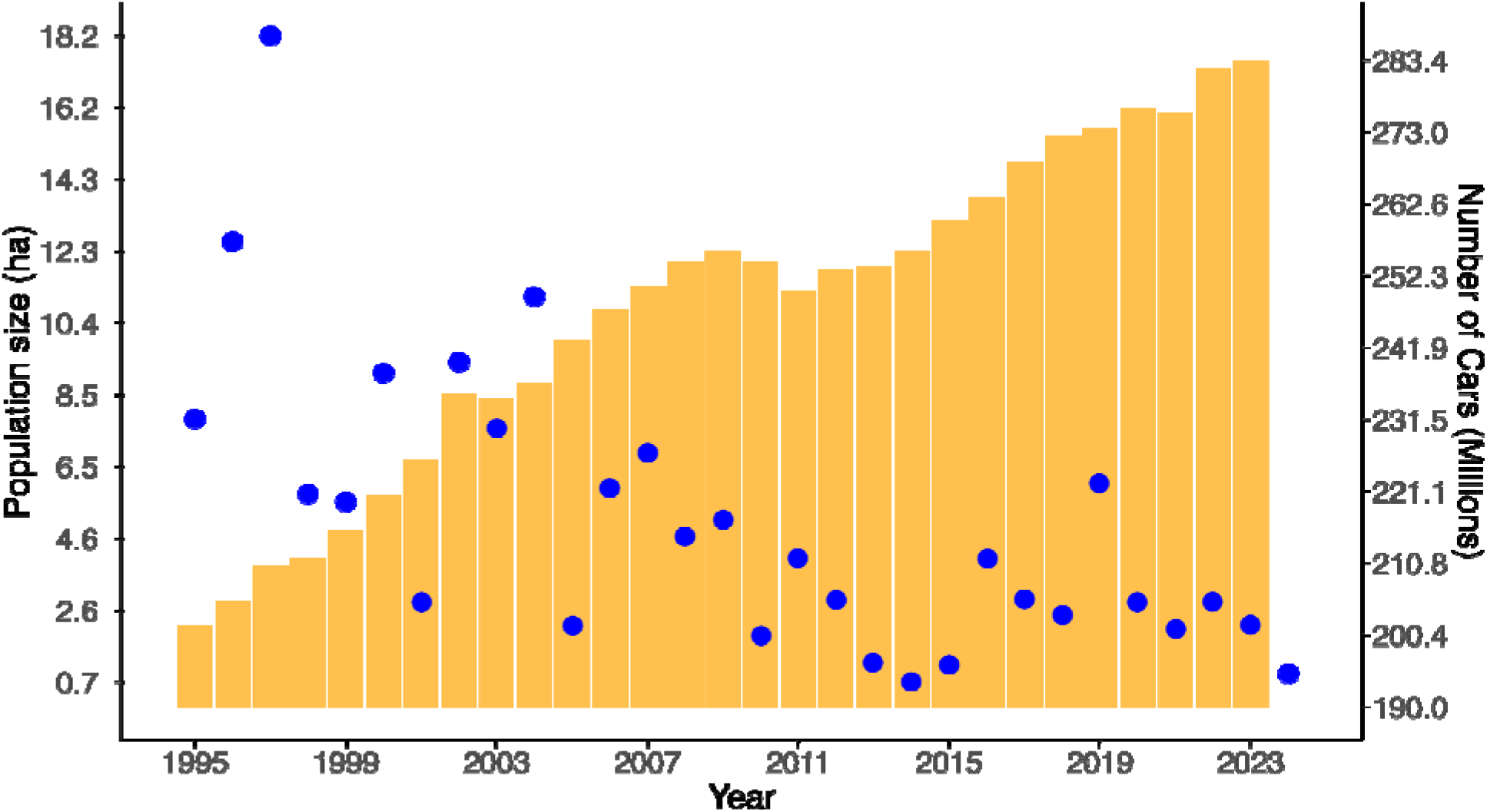
Population size of overwintering migratory monarch butterflies in Mexico in a given year (blue dots) (1) and the number of cars (orange bars) registered in the USA in the previous year (38, 39).

Research to date suggests that habitat loss, specifically loss of breeding host plants, has likely played a large role in the monarch population decline (3–5). Monarchs lay their eggs on milkweed plants, where the larvae grow and develop into adults. Between the mid-1990s and the first decade of the current century the use of the herbicide glyphosate became widespread throughout the central United States and Canada, with huge impacts on milkweed distribution.

For example, Hartzler (6) estimated a 90% decline in milkweed in Iowa corn and soybean fields between 1999 and 2009. Thus, it has been suggested that large-scale milkweed restoration is the most important measure for protection and recovery of the migratory monarch population (4, 7).

Although large-scale milkweed restoration is likely a necessary condition for monarch recovery, it may not be a sufficient condition for recovery if other impacts on monarchs limit the population. One such possible impact is road mortality. Over recent decades, traffic volumes on roads across North America have risen sharply (Fig. 1), and monarchs are frequently-observed roadkill casualties (8–10). This leads to the question: is the road mortality rate high enough that recovery measures aimed only at increasing the distribution of milkweed may be insufficient to recover the population? Previous studies at local or regional scales have suggested that roadkill poses a significant threat to migratory monarchs (11–13), but the potential impact of roadkill has not yet been evaluated on a continental scale. The first step in such an evaluation, and the goal of our study, is to estimate the proportion of migrating monarchs that are killed crossing roads during the fall migration.

We estimated the likelihood of migrating monarchs being killed on a road during fall migration from their breeding range in North America to their overwintering areas in Mexico. We did this for monarchs beginning anywhere in their breeding range, using best estimates of their pre-migration (late summer) distribution, traffic volumes for all roads in North America, monarch flying speed, and flying time spent at traffic height. For the pre-migration distribution we used the map produced by Momeni-Dehaghi et al. (14) based on community science data. For traffic volumes and roads, we used data from OpenStreetMap (OSM) and the US Department of Transportation (15, 16), to produce an estimated average traffic volume for different road types.

For flying speed, we used the average speed of monarchs reported by Knight et al. (17) and for flying time spent at traffic height we polled monarch researchers for expert opinion. We then combined this information in a simulation model to estimate the average probability of a monarch dying due to roadkill during fall migration. To achieve this, we assigned 100 million virtual butterflies to pixels in the pre-migration monarch range, in proportion to the relative abundance estimates for each pixel. Each butterfly’s survival probability was then the product of the probability of surviving all road crossings along the shortest path connecting its starting pixel to the overwintering habitat in Mexico. For each individual road on the path, the survival probability was the product of the estimated proportion of flying time spent at traffic height and survival probability given monarch flying speed and the traffic volume, estimated using the approach in Hels and Buchwald (18).

## Methods

### Premigration distribution

Momeni-Dehaghi et al. (14) estimated the pre-migration distribution of eastern monarchs in 100 × 100 km pixels, using community science data while accounting for sampling bias. In this study, we used this map to create 100 million virtual monarchs, distributed across North America. Virtual monarchs began their fall migration from the centroid of these 100 × 100 km pixels.

### Migration routes

We aimed for a conservatively low estimate of the number of roads crossed by monarchs during their fall migration. Thus, we defined the migration routes as the shortest path from the centroid of pre-migration distribution pixels to the overwintering habitats in Mexico. To map the shortest path for each pixel, we used the Least Cost Path function from the QGIS software (www.qgis.org), considering latitude as the cost. This approach ensures that moving southward (towards the overwintering habitat) always incurs a lower cost compared to moving eastward, westward, or northward. It has been observed that monarchs typically avoid crossing high mountains during their migration. Thus, in our least cost path analysis, we assigned a high cost value (10,000 + latitude) to each pixel of mountainous regions (36).

### Roads and traffic data

As we had traffic data for a subset of all roads in North America, we used traffic data and associated road class to project traffic level on all roads except link roads (see below). We obtained traffic data for various states in the USA for 2017 from the US Department of Transportation. For road classes, to ensure consistency across three countries, we used OpenStreetMap (OSM) road data (16). We selected motorways, primary, residential, secondary, tertiary, trunk, and unclassified roads for our study, excluding all link roads due to insufficient traffic rate data for these types of roads. Upon visually inspecting the data, we observed instances, such as at toll stations and airports, where instead of representing each road with a single line for each direction, each lane was represented by its own line (e.g. Fig. S1). To avoid overestimating the number of roads each monarch butterfly crosses during its migration, we manually adjusted the original dataset by eliminating these excess lines, ensuring only one line per direction was retained. Following this, we transferred traffic rates from traffic layers into corrected OSM layers using the methods described in detail in Supplementary Material S1.

### Flying speed

Given that the flying speed of migratory monarchs is influenced by various factors, including climate, we used the average flying speed of 3.3 m/s during their fall migration, from Knight et al. (17).

### Width of vehicles

The width of the killing zone, defined in our study as the width of the vehicles, is one of the factors affecting the calculated roadkill probability. Wider vehicles increase roadkill probability, as species require more time or higher speed to pass the danger zone. We used the conservative approach of assuming all vehicles were 1.76 m wide (37), which is the average width of passenger vehicles.

### Percent of flying time at car level

To our knowledge, there are currently no data on the proportion of flying time that monarch butterflies spend near the ground during their fall migration. To estimate this variable for our analysis, we used expert opinion. We contacted 55 experts with a research history in monarch butterflies, eliciting their estimates regarding the percentage of flying time that migrating monarchs fly close to the ground (i.e. car height). Their responses were used to adjust our estimates of roadkill probability for monarchs crossing roads at the height of cars.

### Survival/roadkill probability

To estimate the probability of survival for monarchs crossing each road, we used the equation developed by Hels and Buchwald (18):

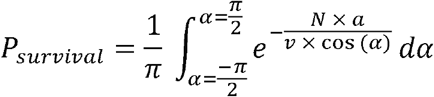

This formula estimates the survival probability based on the traffic rate (N), the width of the killing zone (a), the speed of the animal (v), and the angle of road crossing (α). We made the conservative assumption that monarchs cross all roads perpendicularly (α = 0), which simplifies the equation to:

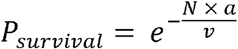

We then adjusted the roadkill rate for each road based on the proportion of flying time that monarchs fly at car level, as elicited from experts’ opinions:

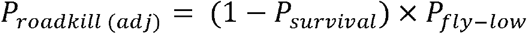

We summarized the survival rate for each migration route using the adjusted roadkill rate:

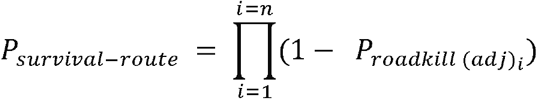

Finally, we calculated the average survival rate across all migration routes by applying the calculations above to each of the 100 million virtual monarchs, and determining the proportion that survived.

### Sensitivity analysis

Given the uncertainties surrounding some of the variables used to estimate survival probability, we conducted a sensitivity analysis. This involved assigning different values to the monarchs’ flying speed and the percentage of flying time at car level. For flying speed, we used 20 equally spaced values ranging from a minimum of 0.1 m/s to twice the standard deviation of flying speeds reported by Knight et al. (17). Similarly, for the percentage of flying time at car level, we used 20 equally spaced values derived from the range of values elicited from experts (Fig. 2).

**Figure 2:**
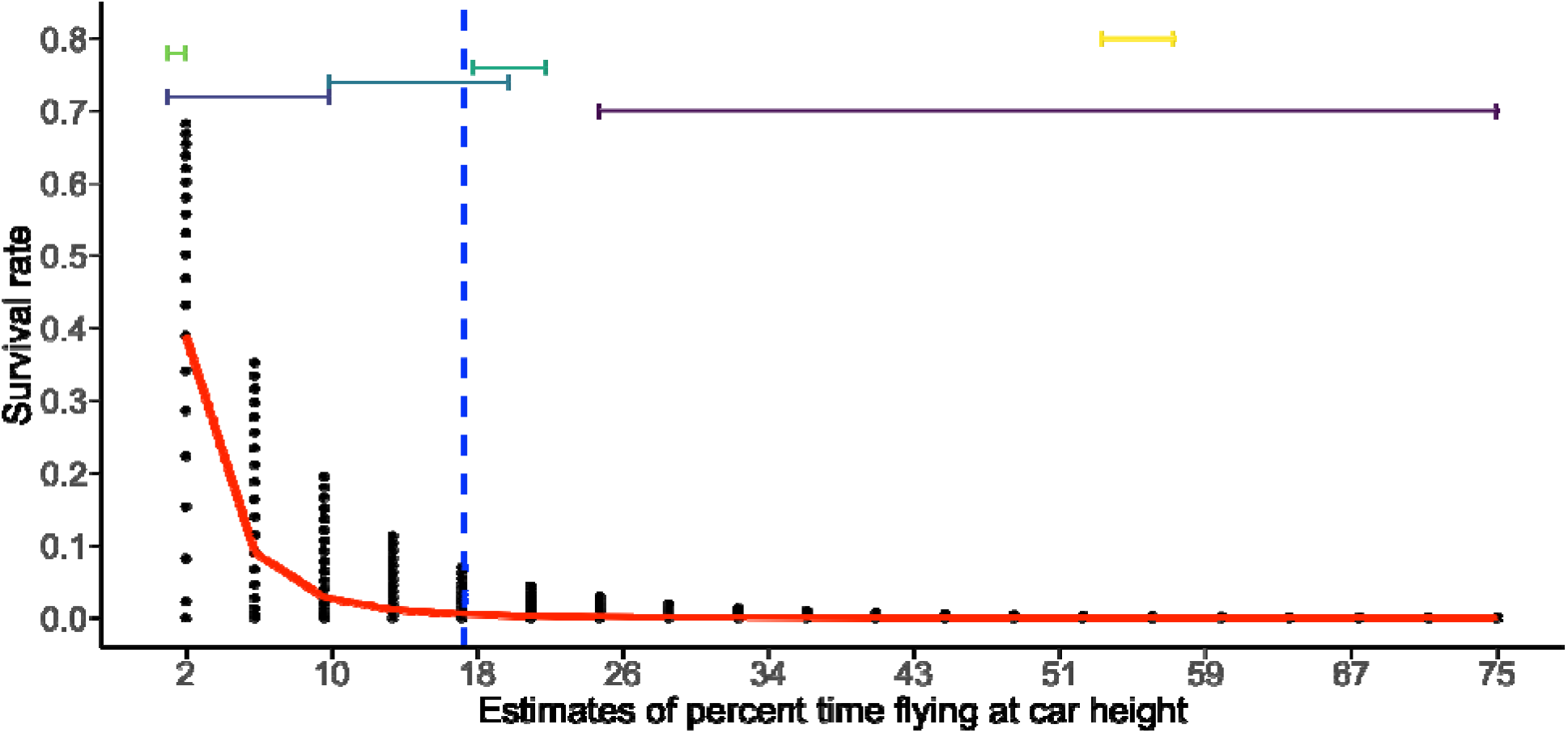
Probability of the average monarch butterfly surviving, i.e. not being killed on a road, during its fall migration to the overwintering grounds in Mexico, for each of 400 different combinations of monarch flying speeds and proportion of flying time at car level. Coloured bars denote the experts’ opinions on the percentage of time that monarchs spend flying at car level during their migration (different bars correspond to different experts). The vertical dashed line shows the median of these estimates.

## Results

Our analysis suggests that, on average, a migratory monarch will migrate 3,283 km (95% CI [3,204, 3,364]) and cross 3,089 roads (95% CI [2,983, 3,222]) during the fall migration if they use the shortest path and are not killed en route. Although we contacted 55 experts for estimates on the percentage of time that monarchs spend flying at traffic height during their fall migration, only six provided a numerical estimate. These estimates varied widely, from 2% to 75% of the flying time (median = 17.5%). Given this range, our analysis suggests that roadkill mortality rate is between 61% and 99.99% (Fig. 2).

## Discussion

Our results suggest that roadkill during the fall migration is likely a crucial factor in the ongoing decline of the migratory monarch population. Previous studies have shown that other factors are also important, especially reduced milkweed abundance leading to reduced breeding possibilities. However, the magnitude of the road mortality rate is such that it calls into question whether recovery measures aimed only at increasing the distribution of milkweed are likely to recover the population if they are not accompanied by measures to reduce roadkill rate.

Although the range of estimates for roadkill was high, the true mortality rate is probably not at either extreme. Our low estimate (61%) may be an underestimate for several reasons. First, we only estimated roadkill rates during the fall migration. Roadkill also occurs during spring migration and the spring-summer breeding period. Mortality rates in spring and summer may be even higher, as the butterflies likely spend a larger proportion of their flying time at car height when they are actively looking for breeding sites. Second, we deliberately made several conservative choices and assumptions in our calculations: (i) we excluded roadkill on link roads due to insufficient traffic data; (ii) we did not adjust traffic estimates for the facts that monarchs are active during daytime when traffic rates are highest (19) and that fall migration occurs from August to October, the time of year when traffic rates are highest (19); (iii) we minimized the likelihood of a monarch crossing a road at traffic height being killed, by assuming a passenger car cross-section and ignoring the higher roadkill risk from larger vehicles such as trucks or buses; (iv) we disregarded indirect mortalities caused by the airflow from passing vehicles (13); (v) we minimized the time a monarch spends on the road by using a high estimate of monarch flying speed (3.3 m/s from Knight et al.(17), as opposed to 2.1 m/s from Matter et al. (20)) and by assuming that monarchs cross roads perpendicularly; and (vi) we minimized the number of roads crossed per monarch by assuming that they follow the shortest route from their starting point to their overwintering habitats in Mexico. On the other hand, our high estimate (99.99%) is likely an overestimate, because this mortality rate suggests imminent extinction.

While ours is the first study to attempt to estimate monarch roadkill probability for fall migration over the whole migratory population, we can compare our estimates to smaller-scale studies directly estimating monarch roadkill numbers over specific road segments. There are three such studies (11–13), but the Mora-Alvarez et al. (13) study was conducted on only very high-traffic roads and therefore is not representative of an average roadkill rate for the continent. In contrast, Kantola et al. (11) and McKenna et al. (12) surveyed roadkilled monarchs across multiple road types in Texas and Illinois, USA, respectively. We extracted the average number of monarchs killed per 100 m of road from each study and calculated the equivalent value from our simulations (Supplementary Materials, Table S2). This comparison indicates that our minimum estimate of 61% mortality, assuming migrating monarchs spend 2% of flying time at car level, is likely an underestimate. This is further supported by the fact that direct roadkill counts, such as those in Kantola et al. (11) and McKenna et al. (12), are noted by the authors as being underestimates due to incomplete detection and low carcass persistence of road-killed butterflies. For example, on moderate-traffic roads, Skórka (21) found that 70% of road-killed butterflies on pavement and 25% of those in verges had disappeared within 24 hours, and within 48 hours almost all had disappeared (99% and 92%, respectively). The number of mortalities per 100 m predicted using our median estimate of 17% of flying time at car level is higher than the roadkill counts in Kantola et al. (11) and McKenna et al. (12) (Supplementary Materials, Table S2), but it is possible that their counts would approach our median estimate if they were adjusted for detectability and carcass persistence.

The large impact of road mortality on the monarch population size implied by our results is indirectly evidenced in previous studies. For example, both Flockhart et al. (22) and Oberhauser et al. (23) found that mitigation efforts to increase the monarch population size were less effective in northern areas than in southern areas. This observation might be due to the lower migration survival rates for monarchs from northern regions, which must cross more roads on their route, thereby reducing the perceived effectiveness of mitigation efforts in the north. In addition, Davis et al. (24) suggest a rapid decline in eastern monarch roost sizes during fall migration, with the rate of decline increasing at lower latitudes. Although roadkill was not identified as a possible cause of this pattern, it is consistent with attrition due to traffic mortality as monarchs migrate southward.

Some density-dependent compensation for fall-migration roadkill may occur during the multi-generational spring-summer breeding season. Negative density-dependence has been shown in monarchs, due to lower larval survival rates on milkweed plants with higher larval densities (25, 26). At even our lowest roadkill estimate of 61%, density-dependent compensation would require each fall-migrating female to replace itself with 5 migrating adults the next fall. However, it is very likely that our lowest roadkill estimate is an underestimate (see above). If our median estimate of 99.5% roadkill is closer to being accurate, this would mean each female would need to replace itself with up to 400 migrating adults. This level of compensation is unlikely, particularly given that roadkill also occurs during the spring and summer.

There are three important unknowns that affect our roadkill estimates. First, the time monarchs spend at car level is highly uncertain and has a large effect on the estimate (Fig. 2). During fall migration, monarchs typically fly high to use tailwind power and reduce energy consumption. However, it has been observed that in the presence of headwinds (27), as well as when nectaring and roosting, monarchs fly much lower, thereby increasing their risk of roadkill. For example, Mora Alvarez et al. (13) observed monarchs searching for suitable roosting places at low elevation next to a highway, increasing their risk of roadkill at a place that is already identified as a roadkill hotspot for monarchs.

Second, there is no information about whether monarchs behaviourally avoid roads or traffic. For example, if they selected migration routes with fewer intersections with roads (i.e., road avoidance behaviour) it could reduce their roadkill risk. However, as mentioned above, by assuming straight-line paths to the overwintering area and perpendicular road crossing, we likely underestimated the area of roads crossed by monarchs. In any case, unlike for flightless insects (28), to date there is no evidence for road avoidance behaviour in monarchs. Indeed, the only study on this topic suggests that some monarchs become accustomed to road-related stressors, such as traffic noise (29), potentially increasing the roadkill risk.

Third, road mortality could cause directional selection pressure on monarch butterflies, favoring individuals that tend to fly at higher altitudes and are therefore less likely to collide with vehicles. Over successive generations, this might lead to a shift in the population’s flight behavior or morphology if such traits are heritable. However, it remains unclear whether this potential adaptation could occur rapidly enough, or to a sufficient extent, to offset the increasing risks posed by expanding road networks and higher traffic rate. Therefore, the capacity for natural selection to mitigate the long-term effects of road mortality on monarch populations is uncertain.

We focused on a specific population, migratory monarchs, as it is the subject of international conservation agreements. However, it is likely that roadkill is an under-appreciated threat to many flying insect species (12, 28, 30–32). For example, Baxter-Gilbert et al. (33) extrapolated from field samples to estimate that, in just one season, North America might witness the roadkill of hundreds of billions of insects, mostly bees, wasps, butterflies, moths, and flies.

A high monarch roadkill rate will make the recovery of the migratory monarch more challenging than if the only major threat were declines in milkweed, the monarch’s breeding host plant. For example, roadsides have been recommended as prime locations for milkweed planting (7) and in some regions governments have begun to apply this recommendation. However, such plantings could inadvertently exacerbate the decline of the monarch population (34, 35) by increasing the time they spend at lower elevations near highways, especially in those regions where milkweed has not senesced prior to the migration period. Therefore, we suggest that recovery efforts should carefully consider the likely impact of roadkill on persistence of the migratory monarch butterfly.

## Acknowledgements

We thank Risa Sargent for her comments and sharing her ideas throughout this study. We thank Sahebeh Karimi and Justin Kreller for their scientific consultation. We thank Gurobi Optimization for providing a free academic license. We thank all the scientists who responded to our question and expressed their best estimates of the percent of time monarchs fly close to the ground. Road data copyrighted OpenStreetMap contributors and available from https://www.openstreetmap.org.

## Author contributions

Conceptualization: IMD, JRB, LF, GWM, TR

Methodology: IMD, JRB, LF, GWM, TR, JOH

Investigation: IMD, JOH, GWM

Visualization: IMD

Funding acquisition: JRB, LF

Project administration: JRB, LF

Supervision: JRB, LF, GWM, TR

Writing – original draft: IMD, JOH

Writing – review & editing: IMD, JRB, LF, GWM, TR, JOH

## Data and materials availability

Traffic data are freely available to download using this link: https://www.fhwa.dot.gov/policyinformation/hpms/shapefiles_2017.cfm. The OSM data are freely available to download using this link: https://download.geofabrik.de/. The R codes we used for analysing the data and visualization are available to download from https://osf.io/jfmnc/?view_only=7cd8fdecd6ee49269e5c2e2818269a2b.

## References

1. E. Rendón-Salinas, A. Fernández-Islas, M. Cruz-Piña, G. Mondragón-Contreras, A. Martínez-Pacheco, “Area of forest occupied by the colonies of monarch butterflies in Mexico during the 2023-2024 overwintering period” (WWF México, 2024).

2. J. E. Diffendorfer, et al., The benefits of big-team science for conservation: Lessons learned from trinational monarch butterfly collaborations. Front. Environ. Sci. 11, 1079025 (2023).

3. J. M. Pleasants, K. S. Oberhauser, Milkweed loss in agricultural fields because of herbicide use: effect on the monarch butterfly population. Insect Conserv. Divers. 6, 135–144 (2012).

4. C. Stenoien, et al., Monarchs in decline: a collateral landscapelJlevel effect of modern agriculture. Insect Sci. 25, 528–541 (2018).

5. W. E. Thogmartin, et al., Monarch butterfly population decline in North America: Identifying the threatening processes. R. Soc. Open Sci. 4, 170760 (2017).

6. R. G. Hartzler, Reduction in common milkweed (Asclepias syriaca) occurrence in Iowa cropland from 1999 to 2009. Crop Prot. 29, 1542–1544 (2010).

7. W. E. Thogmartin, et al., Restoring monarch butterfly habitat in the Midwestern US: ‘all hands on deck.’ Environ. Res. Lett. 12, 074005 (2017).

8. J. Laidman, Monarchs mass for annual trek. Toledo Bl. 33 (1999).

9. F. W. Strong, Outdoors with Fred, Tagging butterflies … yet. Vic. Advocate 38 (1970).

10. M. Ugarte, Monarch butterflies bulk up for migration. Sarasota Her.-Trib. XB (2003).

11. T. Kantola, J. L. Tracy, K. A. Baum, M. A. Quinn, R. N. Coulson, Spatial risk assessment of eastern monarch butterfly road mortality during autumn migration within the southern corridor. Biol. Conserv. 231, 150–160 (2019).

12. D. D. McKenna, K. M. McKenna, S. B. Malcom, M. R. Berenbaum, Mortality of lepidoptera along roadways in central Illinois. J. Lepidopterists Soc. 55, 63–68 (2001).

13. B. X. Mora Alvarez, R. Carrera-Treviño, K. A. Hobson, Mortality of monarch butterflies (Danaus plexippus) at two highway crossing “hotspots” during autumn migration in Northeast Mexico. Front. Ecol. Evol. 7, 273 (2019).

14. I. Momeni-Dehaghi, J. R. Bennett, G. W. Mitchell, T. Rytwinski, L. Fahrig, Mapping the premigration distribution of eastern Monarch butterflies using community science data. Ecol. Evol. n/a (2021).

15. FHWA, Highway Performance Monitoring System (HPMS). Deposited 2017.

16. Geofabrik GmbH, OpenStreetMap Data Extracts. Deposited 2023.

17. S. M. Knight, G. M. Pitman, D. T. T. Flockhart, D. R. Norris, Radio-tracking reveals how wind and temperature influence the pace of daytime insect migration. Biol. Lett. 15, 20190327 (2019).

18. T. Hels, E. Buchwald, The effect of road kills on amphibian populations. Biol. Conserv. 99, 331–340 (2001).

19. M. E. Hallenbeck, M. Rice, B. Smith, C. Cornell-Martinez, J. Wilkinson, “Vehicle volume distributions by classifications” (United States. Federal Highway Administration, 1997).

20. S. F. Matter, A. F. Parlin, O. R. Taylor, J. A. Rich, P. A. Guerra, Meteorological Conditions and Flight Speed during Observed Eastern Monarch Fall Migration Events. J. Lepidopterists Soc. 78 (2024).

21. P. Skórka, The detectability and persistence of road-killed butterflies: An experimental study. Biol. Conserv. 200, 36–43 (2016).

22. D. T. T. Flockhart, J.-B. Pichancourt, D. R. Norris, T. G. Martin, Unravelling the annual cycle in a migratory animal: breeding-season habitat loss drives population declines of monarch butterflies. J. Anim. Ecol. 84, 155–165 (2015).

23. K. S. Oberhauser, et al., A trans-national monarch butterfly population model and implications for regional conservation priorities. Ecol. Entomol. 42, 51–60 (2017).

24. A. K. Davis, J. R. Croy, W. E. Snyder, Dramatic recent declines in the size of monarch butterfly (Danaus plexippus) roosts during fall migration. Proc. Natl. Acad. Sci. 121, e2410410121 (2024).

25. D. T. T. Flockhart, T. G. Martin, D. R. Norris, Experimental examination of intraspecific density-dependent competition during the breeding period in monarch butterflies (Danaus plexippus). PLoS ONE 7, e45080 (2012).

26. L. Marini, M. P. Zalucki, Density-dependence in the declining population of the monarch butterfly. Sci. Rep. 7, 13957 (2017).

27. D. L. Gibo, M. J. Pallett, Soaring flight of monarch butterflies, Danaus plexippus (Lepidoptera: Danaidae), during the late summer migration in southern Ontario. Can. J. Zool. 57, 1393–1401 (1979).

28. P. T. Muñoz, F. P. Torres, A. G. Megías, Effects of roads on insects: a review. Biodivers. Conserv. 24, 659–682 (2015).

29. A. K. Davis, H. Schroeder, I. Yeager, J. Pearce, Effects of simulated highway noise on heart rates of larval monarch butterflies, Danaus plexippuslJ: implications for roadside habitat suitability. Biol. Lett. 14, 20180018 (2018).

30. A. E. Martin, S. L. Graham, M. Henry, E. Pervin, L. Fahrig, Flying insect abundance declines with increasing road traffic. Insect Conserv. Divers. 11, 608–613 (2018).

31. R. S. P. Rao, M. K. S. Girish, Road kills: Assessing insect casualties using flagship taxon. Curr. Sci. 92, 830–837 (2007).

32. D. A. Soluk, D. S. Zercher, A. M. Worthington, Influence of roadways on patterns of mortality and flight behavior of adult dragonflies near wetland areas. Biol. Conserv. 144, 1638–1643 (2011).

33. J. H. Baxter-Gilbert, J. L. Riley, C. J. H. Neufeld, J. D. Litzgus, D. Lesbarrères, Road mortality potentially responsible for billions of pollinating insect deaths annually. J. Insect Conserv. 19, 1029–1035 (2015).

34. W. Keilsohn, D. L. Narango, D. W. Tallamy, Roadside habitat impacts insect traffic mortality. J. Insect Conserv. 22, 183–188 (2018).

35. P. Skórka, M. Lenda, D. Moroń, K. Kalarus, P. Tryjanowski, Factors affecting road mortality and the suitability of road verges for butterflies. Biol. Conserv. 159, 148–157 (2013).

36. UNEP-WCMC, Mountains of the World. 10.34892/dvwq-4j45. Deposited 2002.

37. S. Meyer, Average car size is increasing — will roads still be safe for small cars and pedestrians? (2024). Available at: https://www.thezebra.com/resources/driving/average-car-size/ [Accessed 8 April 2024].

38. Bureau of Transportation Statistics (BTS), Number of U.S. aircraft, vehicles, vessels, and other conveyances. Deposited 4 April 2024.

39. CEIC data, United States number of registered vehicles. (n.d.). Available at: https://www.ceicdata.com/en/indicator/united-states/number-of-registered-vehicles [Accessed 22 April 2024].

