## Supplementary materials for "Roadkill is a crucial factor in the population decline of migratory monarch butterflies"

**S1: Detailed description of transferring traffic data into the OSM road layers**

We developed a dataset to quantify traffic rates for different types of roads across North America. As described in the main text, this dataset was created by merging traffic data (1) with OpenStreetMap road data (hereafter, referred to as OSM data) (2). We needed to merge these two datasets together because (i) traffic dataset did not include data for many roads in our study area, and (ii) OSM data provides an almost complete inventory of roads, featuring a classification system that is relatively consistent across North America (3). Due to the absence of standardized identifiers for reliably merging these datasets (e.g., many roads lacked road names), we developed a method that combines these datasets based on spatial proximity and optimization techniques.

Here, we outline the rationale and steps for the procedure used to merge the traffic and OSM datasets. In broad terms, the procedure entails the following steps: (1) importing and cleaning the OSM and traffic data; (2) subdividing the roads in these datasets into segments and subsequently generating points for each segment; (3) identifying pairs of OSM points that correspond to opposing lanes on two-way roads; (4) matching up OSM points with traffic data points to allocate traffic rate values to the OSM points; (5) addressing discrepancies in the traffic rate values assigned to OSM points; and (6) aggregating the traffic estimates for OSM points into each corresponding OSM segment. These analyses were completed using the *sf* (4), *nngeo* (5), *gurboi* (6), *lwgeom* (7), and *whitebox* (8, 9) *R* packages. Below we provide further details on each of these steps (refer to the Data and materials availability section for implementation details).

**Step 1. Importing and cleaning the OSM and traffic data**

In this step, we aimed to exclude roads unsuitable for our analysis, fix underlying geometry issues in the OSM and traffic data, and standardize road names across both lanes of two-way roads. We first filtered the data to retain only roads classified as 'motorway,' 'primary,' 'residential,' 'secondary,' 'tertiary,' 'trunk,' or 'unclassified.' We excluded all link roads, as preliminary data inspection indicated insufficient traffic rate data for this road type. We then standardized the road names for the two-way roads when just a single lane had an assigned name. This standardization procedure was based on the road reference numbers and nearest neighbor criteria.

**Step 2. Subdividing both datasets into segments and generating points for each segment**

This step was necessary for reconciling differences in the underlying geometries of the two datasets (i.e. OSM and traffic data) and matching traffic dataset with the OSM data based on spatial location. This is because (i) there was no consistent identifier that could be used to match the same road across the two datasets, (ii) the two datasets did not agree perfectly on the location for many roads (e.g., a road in one dataset might be located 5 m away from its counterpart in the other dataset), (iii) a road might be represented using multiple distinct geometries (connecting end-to-end) in one dataset, and represented using a single multi-part geometry in another dataset, and (iv) different lanes of two-way roads might be represented using a single spatial line in one dataset and using multiple spatial lines in another dataset (Fig. S2). In this step, we splitted the roads in both datasets into 500-m line segments and then converted them into 20m evenly-spaced points after removing very short segments (≤ 1m).

**Step 3. Identifying and pairing different lanes (directions) of two-way roads**

This step was important so that we could subsequently correct traffic rates by accounting for differences in the spatial representation of two-way roads (i.e., when one dataset represents a two-way road with a single line geometry and the other represents the same road with a pair of line geometries). This step includes identifying the 50 nearest neighboring OSM points for each OSM point within the specified distance and then refining the pairs of candidate points until we identify pairs of OSM points that likely belong to different lanes of the same two-way road. To match pairs of OSM points that corresponded to the same location on opposite lanes of a two-way road, we used Gurobi software (version 10.0-2) and optimization procedure minimizing the squared Euclidean distance between pairs of points along a road segment. We used squared Euclidean distance because visual inspection of preliminary results revealed that simply using Euclidean distance resulted in incorrect pairings.

**Step 4. matching up OSM points with traffic data points**

We spatially joined the OSM points with the traffic data points. This step was important so that we could transfer traffic rates from the traffic data points to the OSM points. The criteria for joining these data were that a given OSM point could only be joined with a traffic data point if (i) the traffic data point was the closest (among all traffic data points) to the given OSM point, and (ii) the OSM given point was the closest (among all OSM points) to the traffic data point. In other words, this was a strict nearest neighbor join. This step was critical for ensuring that multiple OSM points would not be paired with the same traffic data point (and vice versa), and reducing the chances of incorrectly joining OSM points with the wrong traffic data point. We then used information from the OSM points pairings to improve the joining process and errors associated with two-way roads.

**Step 5. addressing discrepancies in the traffic rate values assigned to OSM points**

This was important to ensure that traffic rates would not be overestimated or underestimated due to differences in the spatial representation of two-way roads between the OSM and traffic rate datasets. Specifically, when considering pairs of OSM points (as outlined in step 3), if both points from a pair were matched to the same traffic data point, we halved the traffic rate assigned to each point. This adjustment was necessary because it indicates that the OSM dataset represented a two-way road with two separate spatial lines, whereas the traffic rate dataset depicted the same road with a single spatial line. Therefore, halving the traffic rate is essential to avoid overestimating the actual traffic volume on the road.

**Step 6. aggregating the traffic rates for OSM points into each corresponding OSM segment**

After correcting the traffic rates for the OSM points, we used these data to calculate traffic rates for the OSM segments. To achieve this, we identified the OSM points that occurred along each segment and calculated the traffic rates for the OSM segment based on the modal value of the OSM points occurring along it.

Thus, for each state, we produced a complete road network dataset that contained traffic rates at least for a part of its roads. By using this process, we were able to import the traffic rate data for 16.9 to 47.7 percent of OSM road segments, depending on different states. Due to the highly computationally intensive nature of this processing, we limited our efforts to the 14 states: Alabama, Arkansas, Connecticut, Delaware, District of Columbia, Florida, Georgia, Indiana, Kansas, Louisiana, Maine, Maryland, Massachusetts, and Michigan. We then determined the average traffic rate for each road type across these states and applied these average rates to all roads of the same type in the study area (Table S1). The entire process spanned 9 days and utilized two computer systems. For reference, these computer systems had the following specifications.

•	Apple M1 Max processor (10 cores) and 64 GB memory.

•	AMD Ryzen 9 5900x processor (12 cores) and 128 GB memory.

**S2: Methodology for Comparing Calculated Roadkill Probabilities with Previous Studies**

**Overview**

Three field studies to date have published estimates of the number of roadkilled monarchs with respect to traffic. Mora-Alverez et al. (10) surveyed roadkilled monarchs in 2018 along highways in the states of Coahuila and Nuevo León, Mexico, Kantola et al. (11) surveyed roadkilled monarchs across three road types in 2016 and 2017 in Texas, United States of America, and McKenna et al. (12) surveyed roadkilled monarchs in 1998 across five road types in the state of Illinois, United States of America. To compare monarch roadkill estimates between these studies and with our study, we had to standardize the roadkill estimates from the field studies by survey effort and the length of the migration period at each location. Given that our study was focused on estimating mortality rates associated with road mortality across the migratory range, we also had to estimate the average number of roadkilled monarchs per 100 m of road for our study given different estimated mortality rates, while controlling for the size of the monarch population in the year of each field study to make estimates comparable.

**Standardizing roadkill estimates**

Mora-Alverez et al. (10) estimated the number of roadkilled monarchs areas along two major highways across the entire migration season, at locations where monarchs are known to cross the highways in large numbers and road mortality is readily observed. Surveys along the highway in Nuevo León used a transect width of 3.4 m, while those in Coahuila used a transect width of 2.9 m. Therefore, we standardized their roadkill estimate from Coahuila using a multiplying factor of 1.7 (i.e., 3.4/2.9). We then divided the total estimated number of roadkilled monarchs by the length of the survey transects along each highway to estimate the number of monarchs killed per km. Last, we divided these estimates by 10 and averaged the numbers from both highways, to estimate the number of roadkilled monarchs per 100 m.

Kantola et al. (11) estimated the number of monarchs killed by vehicle collisions in the “central funnel” through Texas and southern Oklahoma, United States of America across two years, based on data collected along roads in Texas, United States of America. They used a MaxEnt model to estimate the average number of monarchs killed across both years across the three road types they surveyed in the “central funnel”, which they estimated to be 2.11 million monarchs. This accounted for surveying on both sides of a road and medians where present, but the transect width used was 1 m. Therefore, we took their estimate of monarchs killed in the “central funnel” and divided this by the km of each road type in the central funnel. We then divided this value by 10 to get the number of monarchs per 100 m and adjusted it by a factor of 3.4 so it is comparable to the data from Mora-Alverez et al. (10).

McKenna et al. (12) surveyed the number of roadkilled monarchs in Illinois, United States of America across five road types within a single migration season. We extracted the average number of monarchs killed per 100 m per day for each week of the study in Figure 4 in McKenna et al. (12). We then averaged these values and multiplied the average by the length of the study, i.e., 42 days, to derive the cumulative number of monarchs killed per 100 m of road. McKenna et al. (12) further used a survey width of 2 m; therefore, we adjusted the average number of roadkilled monarchs per 100 m per day by a factor of 1.7 to be comparable with Mora-Alverez et al. (10). It is important to note that McKenna et al. (12) likely missed the start of the migration season, given that surveys started early September, but migration begins in mid-August, possibly underestimating roadkill mortality. It is also important to note that all surveys in a week took place over a two-day period, i.e., one road was surveyed on a single day for only part of the day, further underestimating the number of roadkilled monarchs because each road survey did not take place over the entire period per day that monarchs are active.

**Estimating number of roadkilled monarchs from our model**

To estimate the number of roadkilled monarchs per 100 m of road over the migration period in our study, we first estimated the starting fall migration population size for monarchs for a given year. This was accomplished by multiplying the estimated area occupied on the wintering grounds in ha by monarchs in Mexico for the same years during which the field studies took place by an estimate of 21.1 million monarchs per ha from Thogmartin et al. (13). We then divided this by two focal survival rates from our study (please refer to figure 2). Specifically, we used survival rates of 0.05 and 0.39, which represent our estimated survival rates from our roadkill model if monarchs spend 17.5% and 2% of their time flying at car level, respectively (Fig. 2). The value of 17.5% represents the median estimate of time spent flying at car level based on survey responses from five monarch experts and 2% represents the lowest estimate received from a single expert (please refer to Methods). This estimate assumes that 100% of fall migration mortality is the result of vehicle collisions. We had to make this necessary and simplifying assumption given there is no empirical data on other causes of mortality and acknowledge that this will likely results in overestimates of the number of monarchs killed per 100 m of road. Next, we subtracted the estimate of the number of monarch butterflies on the wintering grounds from our estimate of the fall migratory population size to derive the number of monarchs that died during migration. Last, we divided this estimate by the total length of roads in our study area in km (7,879,217 km) and divided this number by 10 to get an estimate of the number of roadkilled monarchs per 100 m.

Table S1: Average traffic rate assigned to each road-type in the study area.

| Road type | Traffic direction | AADT | Road type | one/two way | AADT |
| --- | --- | --- | --- | --- | --- |
| Motorway | Both | 47338.9 | Trunk | Both | 7044.8 |
| Motorway | One | 26893.8 | Trunk | One | 9114.8 |
| Primary | Both | 6067.0 | Unclassified | Both | 910.9 |
| Primary | One | 12815.4 | Unclassified | One | 5160.7 |
| Secondary | Both | 4155.9 | Residential | Both | 1473.9 |
| Secondary | One | 8919.8 | Residential | One | 3001.5 |
| Tertiary | Both | 2952.9 | - | - | - |
| Tertiary | One | 5058.3 | - | - | - |

Table S2. Estimated number of roadkilled monarchs per 100 m of road from three field studies and our roadkill model.

| **Study** | **Monarchs/100 m^‡^** | **Wintering years** | **Location** | **Dates** | **Road type** | **Winter density (ha)** | **AADT** | **Adjusted monarchs per 100 m from this study*^†^** |
| --- | --- | --- | --- | --- | --- | --- | --- | --- |
| Mora Alvarez et al. 2019 | 890.9 | 2018/2019 | Coahuila & Nuevo León, Mexico | Oct 22 – Nov 14 (Coachuila) & Oct 24 – Nov 11 (Nuevo León) | Highway only | 6.05 | 14330 (Nuevo León) & 8,862 (Coahuila) | 292.95 (17.5% of time flying at car level) |
|  |  |  |  |  |  |  |  | 2.54 (2% of time flying at car level) |
| Kantola et al. 2019 | 8.7 | 2016/2017 2017/2018 | Texas, USA | Oct 10 – Nov 07  Oct 03 – Oct 27 | Mix (n = 3 types) | 2.70 | Not reported | 130.74 (17.5% of time flying at car level) |
|  |  |  |  |  |  |  |  | 1.13 (2% of time flying at car level ) |
| Mckenna et al. 2001 | 4.3 | 1998/1999 | Illinois, USA | Sep 09 – Oct 19 | Mix (n=5 types) | 5.56 | 7546**^+^** | 269.23 (17.5% of time flying at car level) |
|  |  |  |  |  |  |  |  | 2.33 (2% of time flying at car level) |
| This study (17.5% of time flying at car level)* | 43.6 | 2023/2024 | North America | Aug 15 – Nov 01 (?) | Mix (n=7 types) | 0.90 | 10064**^+^** | NA |
| This study (2% of time flying at car level )* | 3.8 | 2023/2024 | North America | Aug 15 – Nov 01 (?) | Mix (n=7 types) | 0.90 | 10064**^+^** | NA |

^‡^Estimates based on study duration and standardized to a transect survey width of 3.4 m used by Mora-Alvarez et al. (2019), i.e. the largest transect width used in any of the studies.

^+^Average Annual Average Daily Traffic (AADT) across all road types.

*Average number of roadkilled monarchs based on our model across all road types for the specified amount of time migrating at car level.

^†^Adjusted relative to winter density at time of other study

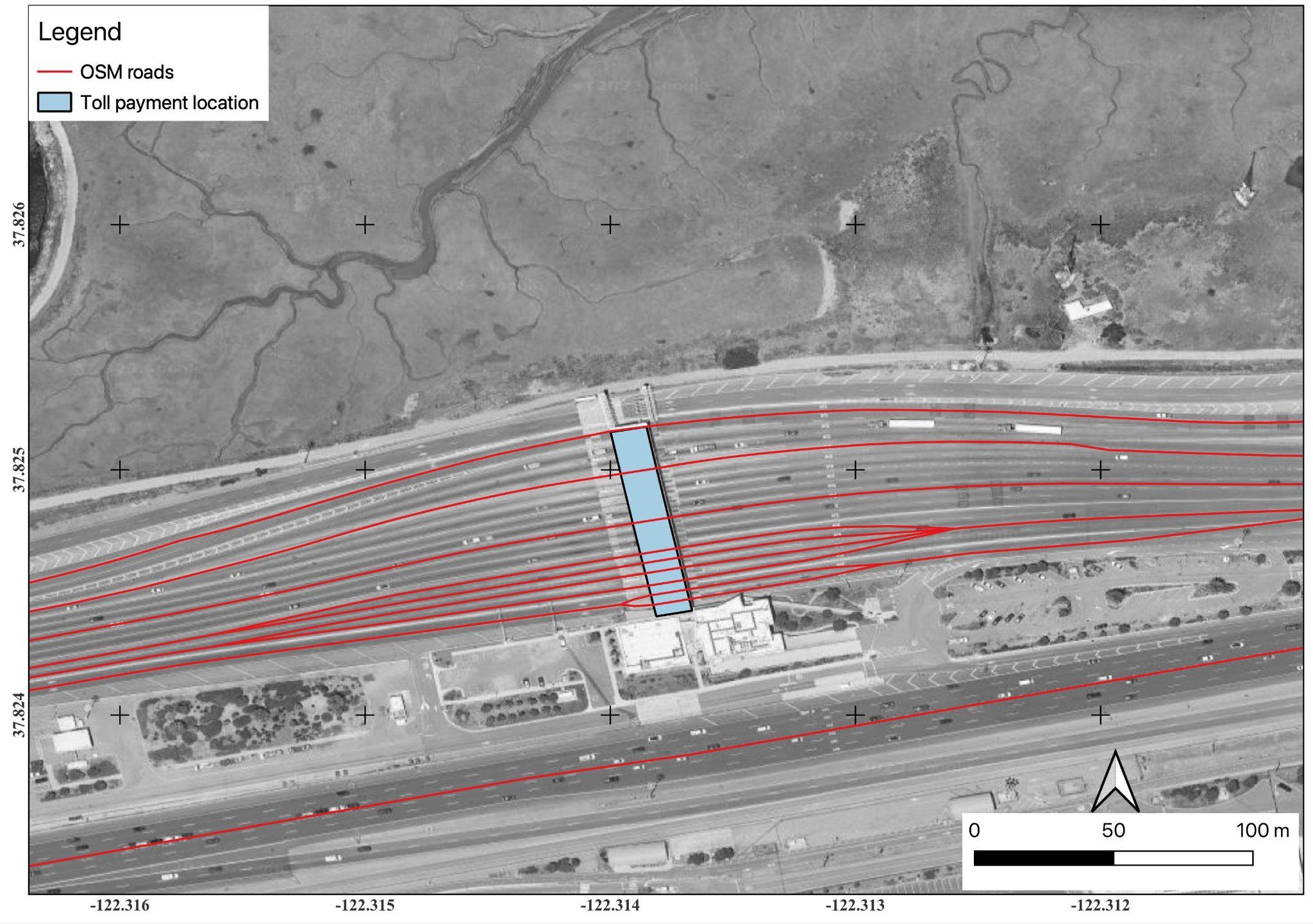

Figure S1 - An example of a toll station on OSM data and how for a single road that has multiple lines in the OSM road layer. Using on-screen inspection of the data, we identified such areas and manually corrected them. The background image is a Google satellite image accessed on 29 May 2023.

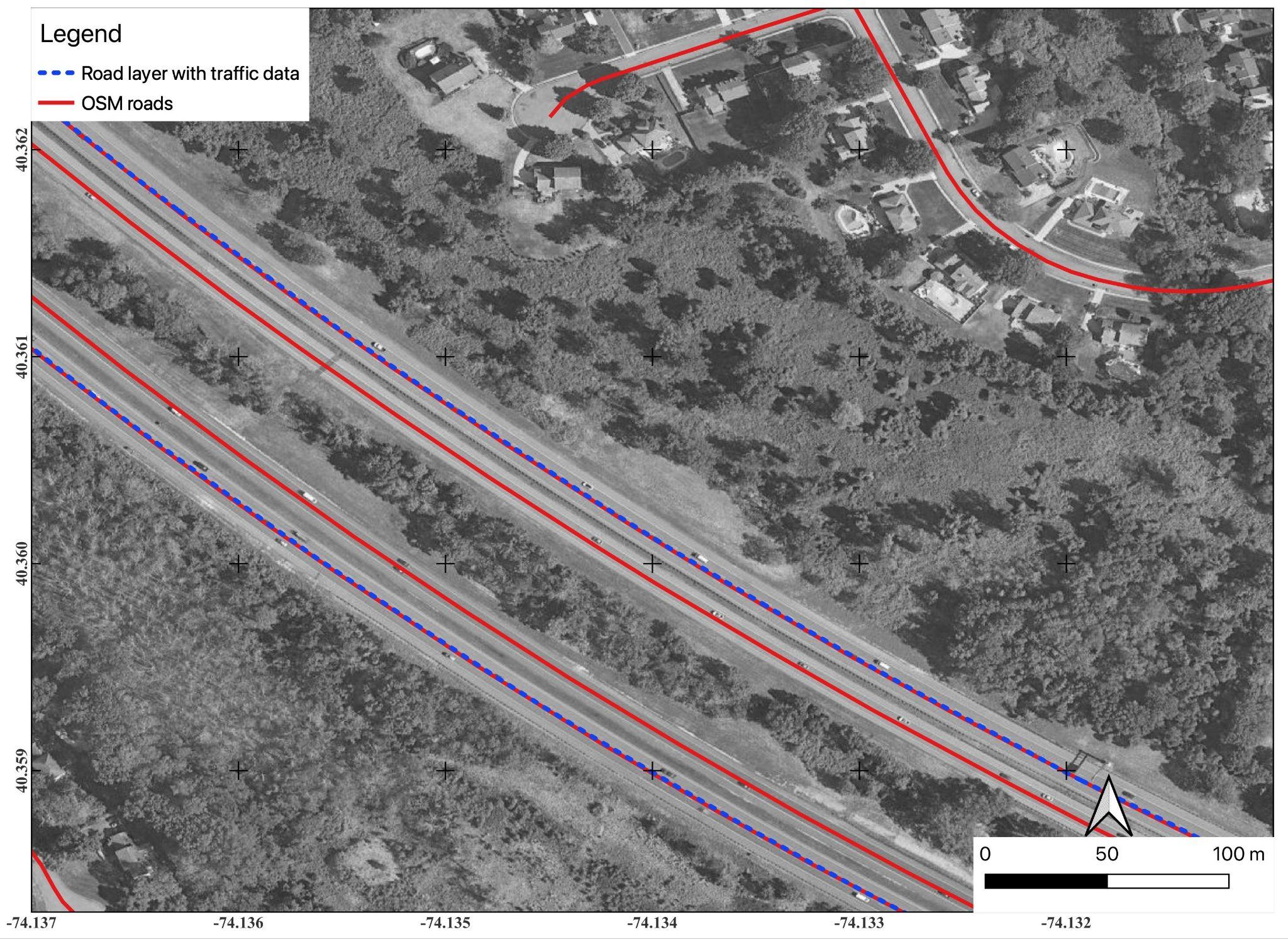

Figure S2 - An example of a road having multiple lines on OSM but a lower number of lines on our road layer containing traffic data. Background image: Google satellite image accessed 29 May 2023.

**References:**

1.	FHWA, Highway Performance Monitoring System (HPMS). Deposited 2017.

2.	Geofabrik GmbH, OpenStreetMap Data Extracts. Deposited 2023.

3.	[OpenStreetMap Wiki contributors, Highway:International equivalence. (2022). Available at: https://wiki.openstreetmap.org/wiki/Highway:International_equivalence [Accessed 9 April 2024].](https://www.zotero.org/google-docs/?smX1Ix)

4.	Pebesma E, Simple features for R: standardized support for spatial vector data. *R J.* **10**, 439–446 (2018).

5.	Dorman M, nngeo: k-nearest neighbor join for spatial data. (2023). Deposited 2023.

6.	Optimization G, LLC, gurobi: Gurobi Optimizer 10.0 interface. (2023). Deposited 2023.

7.	Pebesma E, lwgeom: Bindings to selected ’liblwgeom’ functions for simple features. (2023). Deposited 2023.

8.	J. B. Lindsay, Whitebox GAT: A case study in geomorphometric analysis. *Comput. Geosci.* **95**, 75–84 (2016).

9.	Wu, Q, Brown, A, whitebox: “WhiteboxTools” R Frontend. (2023). Deposited 2023.

10.	B. X. Mora Alvarez, R. Carrera-Treviño, K. A. Hobson, Mortality of monarch butterflies (Danaus plexippus) at two highway crossing “hotspots” during autumn migration in Northeast Mexico. *Front. Ecol. Evol.* **7**, 273 (2019).

11.	T. Kantola, J. L. Tracy, K. A. Baum, M. A. Quinn, R. N. Coulson, Spatial risk assessment of eastern monarch butterfly road mortality during autumn migration within the southern corridor. *Biol. Conserv.* **231**, 150–160 (2019).

12.	D. D. McKenna, K. M. McKenna, S. B. Malcom, M. R. Berenbaum, Mortality of lepidoptera along roadways in central Illinois. *J. Lepidopterists Soc.* **55**, 63–68 (2001).

13.	W. E. Thogmartin, *et al.*, Density estimates of monarch butterflies overwintering in central Mexico. *PeerJ* **5**, e3221 (2017).
